# Prenatal THC exposure alters metabolic gene expression and KEAP1 regulation in the mouse placental labyrinth

**DOI:** 10.64898/2026.11.04.764078

**Authors:** Eleni Sia, Pedro Henrique Gobira, Fenghua Chen, Gülendam Sezer, Heidi Kaastrup Müller, Sâmia Joca, Cristiana Cruceanu, Rodrigo Grassi-Oliveira

**Author notes:** Correspondence: Rodrigo Grassi-Oliveira.

## Abstract

**Introduction:** Prenatal cannabinoid exposure may influence fetal development through changes in placental biology, yet its effects on metabolic gene regulation remain poorly understood. We examined the effects of Δ^9^-tetrahydrocannabinol (THC) and cannabidiol (CBD) on the mouse placental labyrinth zone.

**Methods:** Pregnant C57BL/6JRj mice received THC (5 mg/kg), CBD (5 mg/kg), or vehicle daily by intraperitoneal injection from gestational day 5 to 17.5. Targeted NanoString profiling and pathway analysis were combined with RT-qPCR and protein measurements.

**Results:** THC exposure reduced glutaminase (*Gls*), Kelch-like ECH-associated protein 1 (*Keap1*), and FMS-like tyrosine kinase 3 (*Flt3*) expression (false discovery rate = 0.0904 for each). Pathway analysis identified transcriptional changes in the nine-gene arginine-metabolism set (false discovery rate = 0.0527). No genes or pathways met the significance criteria following CBD exposure. RT-qPCR confirmed lower *Keap1* expression with sexes pooled and identified lower *Gls* and *Flt3* expression in male samples. KEAP1 protein abundance increased despite lower *Keap1* mRNA, while nuclear factor erythroid 2-related factor 2 localization and heme oxygenase 1 (*Hmox1*) expression showed no significant differences.

**Discussion:** These findings identify amino acid metabolic gene expression and KEAP1 regulation as complementary components of the placental response to prenatal THC exposure, connecting transcriptional changes with protein regulation within the placental exchange region.

## Introduction

Cannabis use during pregnancy has increased in several populations [1,2]. Research has largely focused on neurodevelopmental outcomes [3], bringing attention to the placental processes that shape the fetal environment. The placenta coordinates nutrient and gas exchange and endocrine signaling [4]. Its contribution to the placenta–brain axis makes placental responses to cannabinoids relevant to developmental and psychiatric research [5,6].

The placenta expresses a functional endocannabinoid system, including CB1, CB2 and nonclassical cannabinoid targets [7,8]. Δ^9^-Tetrahydrocannabinol (THC) and cannabidiol (CBD), two major constituents of *Cannabis sativa*, engage this biology through different pharmacological actions: THC acts primarily through cannabinoid receptors, whereas CBD has lower affinity and a broader range of targets [9]. Their interactions with placental signaling support examining each compound under the same experimental conditions [10,11].

Gestational THC exposure alters placental vascular development in rodents and nonhuman primates [12–14], and trophoblast models reveal effects on signaling, differentiation and mitochondrial function [15–17]. CBD exposure has been associated with labyrinth vascular defects and reduced fetal growth in rats [18], artery remodeling in mice [19], altered trophoblast respiration and differentiation [20], and changes in human placental tryptophan metabolism and xenobiotic transport [21,22].

Metabolic regulation supports placental growth, differentiation and resource transfer [23,24]. Glutamine contributes to mitochondrial respiration in primary human trophoblasts [25], while endocannabinoid signaling interacts with metabolic processes [26]. Placental adaptation to adverse gestational environments involves changes in nutrient handling and energy metabolism [27], making amino acid metabolic gene regulation a relevant target of prenatal cannabinoid exposure.

The labyrinth zone (LZ) is the principal exchange region of the rodent placenta, containing trophoblast layers and fetal vascular endothelium [28]. Layer-enriched sampling can reveal expression responses obscured in whole-placenta measurements [29]. Labyrinth vascular abnormalities after gestational THC or CBD exposure further support this region as a focus for molecular investigation [14,18].

We combined targeted NanoString profiling and pathway analysis with RT-qPCR and protein measurements to examine metabolic gene regulation in the LZ following gestational THC or CBD exposure. KEAP1 and NRF2-related measurements and analyses by fetal sex connected individual transcript responses with their metabolic and protein-regulatory context.

## Methods

### Animals and ethical approval

Female and male C57BL/6JRj mice were purchased from Janvier Labs (Le Genest-Saint-Isle, France) at five and seven weeks of age, respectively. Animals were housed in standard cages (425 × 266 × 185 mm) at 22 ± 2 °C and 60 ± 5% relative humidity, with food and water available ad libitum. The light–dark cycle was 12:12 h, with lights on at 06:00. Animals underwent a two-week acclimatization period before breeding, and handling was minimized during treatment. All procedures complied with Directive 2010/63/EU and the Danish Act on Animal Experimentation and were approved by the Danish Animal Experiments Inspectorate (Dyreforsøgstilsynet; permit 2023-15-0201-01528).

### Drug preparation

Dronabinol (THC; CAS 1972-08-3, EC 625-153-6) and CBD (batch CBD2202A02, DAC quality) were obtained from Candoro Ethics GmbH (Frankfurt, Germany). THC was prepared according to the manufacturer’s instructions, converting dispensed mass to volume using the reported density (0.95 g/mL). The 5 mg/mL stock, stored at −20 °C, was diluted 1:10 to 0.5 mg/mL. CBD powder was weighed and dissolved directly on treatment days. Drugs were freshly prepared in Kolliphor, ethanol and saline (1:1:18), modified from [30]; the same mixture served as vehicle (VEH). Intraperitoneal dosing was 5 mg/kg, with injection volumes of 0.3–0.5 mL scaled to body weight.

### Gestational exposure and sample collection

After acclimatization, one male and one female shared a cage separated by a barrier for three days. The barrier was removed for overnight mating and reintroduced the next day, designated gestational day (GD) 0. Pregnancy was assessed by vaginal plug and subsequent maternal weight gain when the plug was unclear. Pregnant females were randomly assigned to daily intraperitoneal THC (5 mg/kg), CBD (5 mg/kg), or VEH from GD5 to GD17.5 (Fig. 1A), beginning after implantation [28]. Four hours after the final injection, dams were euthanized by cervical dislocation. Fetuses and placentas were collected, fetal tails retained for sex determination, and whole placentas snap-frozen and stored at −80 °C. NanoString samples were randomly selected from four dams per group, comprising six male-associated and six female-associated placentas per group.

**Figure 1.**
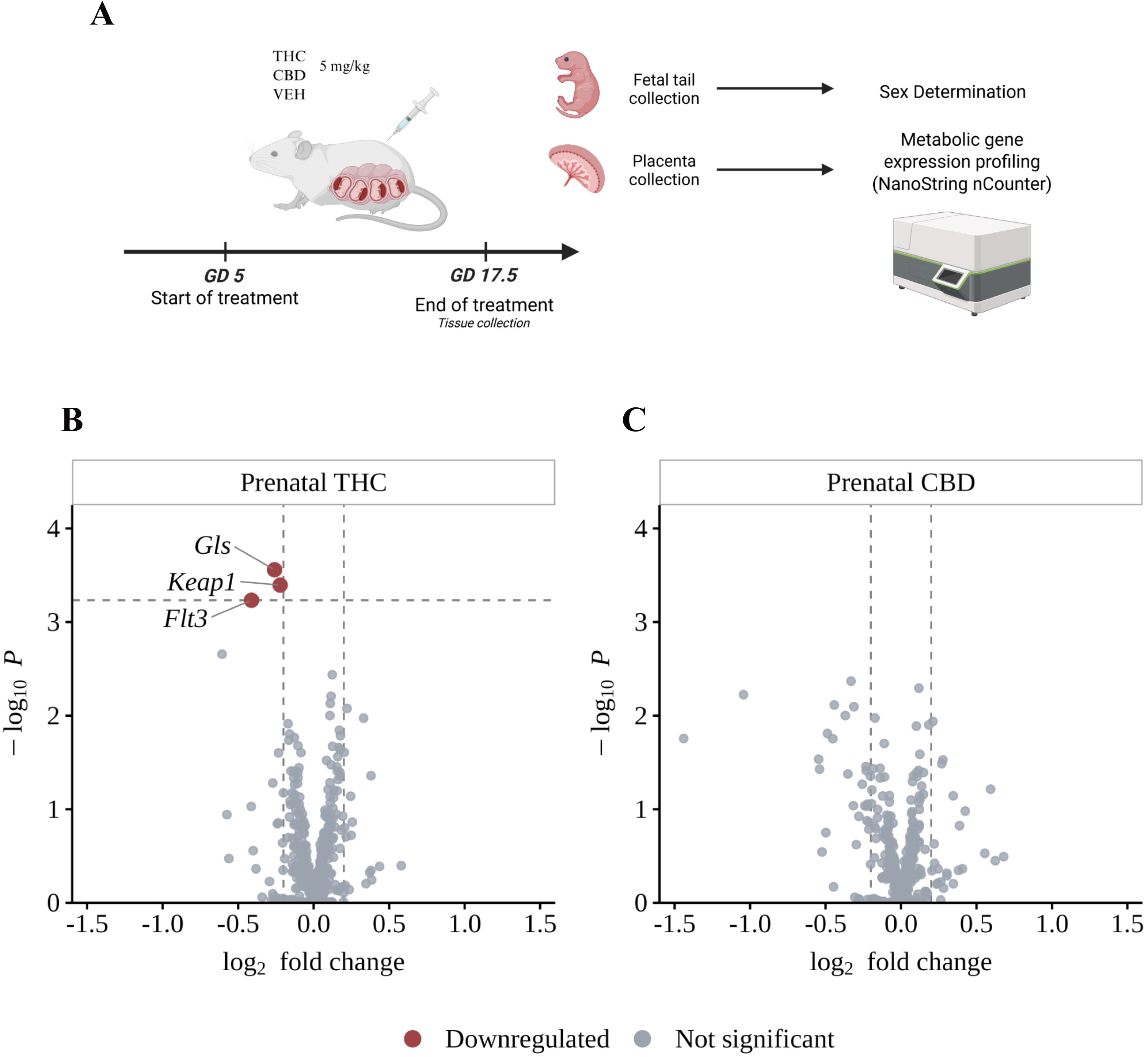
Experimental design and targeted gene expression profiling. (A) Pregnant mice received daily intraperitoneal THC (5 mg/kg), CBD (5 mg/kg), or VEH from GD5 to GD17.5. Placentas and fetal tails were collected four hours after the final injection. Fetal sex was determined by PCR, and LZ tissue was profiled using the nCounter Mouse Metabolic Pathways Panel. The sample comprised 12 placentas per treatment group, six associated with male fetuses and six with female fetuses, selected from four dams per group. (B,C) Volcano plots for THC versus VEH and CBD versus VEH, respectively, based on 463 genes retained at ≥50 raw counts in at least 18/36 samples. Differentially expressed genes were defined by FDR ≤0.10 and |log_2_ fold change| ≥0.2. *Gls*, *Keap1* and *Flt3* were downregulated in the THC-versus-VEH comparison, each with FDR = 0.0904. No genes met both criteria in the CBD-versus-VEH comparison.

### Fetal sex determination

Fetal sex was determined by *Sry* and *Rapsn* PCR with modifications to Behringer et al. [31]. Tail tissue (1–2 mm) was lysed overnight at 55 °C in 50 mM Tris-HCl (pH 8.0), 50 mM EDTA, 0.5% SDS and proteinase K. After enzyme inactivation (95 °C, 10 min) and centrifugation (12,000 × g, 5 min), genomic-DNA supernatants were stored at −20 °C. Amplification used SYBR Green Master Mix (2×), followed by electrophoresis (100 V, 30– 50 min) and UV visualization. *Rapsn* (∼590 bp) indicated DNA integrity in both sexes; *Sry* (∼230 bp) identified males. Primers and cycling conditions are in Supplementary Table S2; Supplementary Figure S1 shows a representative gel.

### LZ sampling and NanoString expression profiling

The LZ was dissected from the frozen tissue on ice, identified using a published dissection guidance [32] and distinguished from the adjacent junctional zone by its characteristic tissue coloration. Total RNA was extracted using the RNeasy Mini Kit (Qiagen, Germany). RNA concentration and purity were assessed with a NanoDrop ND-1000 spectrophotometer (Thermo Fisher Scientific, USA). Samples with A260/A280 ratios of 1.8–2.1 and A260/A230 ratios >1.8 were used for profiling.

Expression profiling used the nCounter Mouse Metabolic Pathways Panel (XT-CSO-MMP1-12; Bruker, USA), comprising 748 target genes and 20 housekeeping genes. A total of 30 ng of RNA was used per hybridization reaction. Hybridization was performed in a thermocycler at 65 °C for 22 h. Following hybridization, samples were loaded into nCounter SPRINT cartridges for automated post-hybridization processing and digital counting using the nCounter SPRINT Profiler (NanoString Technologies). The 36 samples were distributed across three cartridges, each containing two male-associated and two female-associated placentas from each treatment group. Raw data were processed using nSolver Analysis Software 4.0 and the nCounter Advanced Analysis module (version 2.0.134). All samples met the quality-control criteria for imaging efficiency (≥75% of fields of view successfully imaged), binding density (0.1–1.8 spots/µm²), positive-control linearity (R² ≥0.95), and limit of detection (0.5 fM positive-control counts exceeding the mean negative-control counts plus two standard deviations). Expression values were normalized to the geometric mean of 17 housekeeping genes selected using geNorm: *Tbc1d10b*, *Fcf1*, *Tlk2*, *Ubb*, *Dhx16*, *Sdha*, *Sap130*, *Dnajc14*, *Tbp*, *Cog7*, *Oaz1*, *Mrps5*, *Edc3*, *Usp39*, *Stk11ip*, *Agk*, and *Polr2a*. To exclude low-abundance transcripts, target genes were retained for differential-expression analysis if they had at least 50 raw counts in at least 18 of the 36 samples, yielding 463 genes.

Differential expression was assessed in R (version 4.5.2) using the original gene-wise modeling routines implemented in nCounter Advanced Analysis. Models included treatment group as the predictor and fetal sex as a covariate, with VEH as the reference group. P-values were adjusted using the Benjamini–Hochberg procedure across the 463 genes within each treatment-versus-VEH comparison. Differentially expressed genes were defined by an absolute log_2_ fold change ≥0.2 and an FDR ≤0.10. Plots were generated using ggplot2 in R.

### Pathway analysis

Pathway-level transcriptional changes were evaluated using self-contained rotation gene-set testing with mroast in limma (version 3.64.3) [33,34]. The analysis used housekeeping-normalized log_2_ expression values for the same 463 endogenous genes retained at ≥50 raw counts in at least 18 of the 36 samples. Gene sets were defined using the NanoString mouse-panel annotations associated with the study codeset. The primary analysis included 34 metabolic pathways, retaining all eligible genes assigned to each pathway irrespective of individual gene-level significance. Linear models included treatment group and fetal sex, with separate THC-versus-VEH and CBD-versus-VEH contrasts. Samples were modeled as independent observations. Tests used 9,999 rotations, the mean set statistic and fixed random seeds. The nondirectional Mixed test provided the primary pathway-level inference. P-values were adjusted across the 34 pathways within each contrast using the default mroast Benjamini–Hochberg mid-P-value procedure, with FDR ≤0.10 considered significant.

An extended analysis included all 36 categories in the exported annotation, adding antigen presentation and cytokine and chemokine signaling, with FDR recalculated across that family. Directional statistics were retained as complementary measures. Heatmaps displayed gene-wise standardized expression across all 36 samples, arranged by treatment group and fetal sex. Genes contributing to the pathway-level statistic were ranked by the absolute standardized statistics from the pathway linear model.

### Reverse transcription and quantitative PCR

Selected transcripts identified by NanoString profiling were further assessed in the same RNA samples by RT-qPCR [35]. Complementary DNA was synthesized with SuperScript IV reverse transcriptase (Thermo Fisher Scientific, USA) and random primers. Incubation comprised 23 °C for 10 min, 50 °C for 10 min and 80 °C for 10 min. The resulting cDNA preparation was diluted with nuclease-free water to 2 ng/µL. Amplification used SYBR Green master mix (Thermo Fisher Scientific, USA) on an AriaMx Real-Time PCR System (Agilent Technologies, USA), with reactions run in technical duplicate. Cycling conditions were 95 °C for 3 min, followed by 40 cycles of 95 °C for 10 s, 60 °C for 30 s and 72 °C for 1 min.

Primers were designed using Primer-BLAST, and amplification efficiencies ranged from 90% to 110%. Sequences are listed in Supplementary Table S2. Target expression was normalized to the geometric mean of *Gapdh*, *Rpl13a* and *Ywhaz*, and relative expression was calculated as 2^−ΔCq^. Expression is presented as a fold change relative to the mean of the VEH group, multiplied by 100.

### Whole tissue lysates and subcellular fractionation

Whole LZ lysates were prepared by homogenization in STM buffer and further dilution in 1× RIPA buffer containing Halt Protease and Phosphatase Inhibitor Cocktail (Thermo Fisher Scientific, USA; 1:100). Samples underwent three water-bath sonication cycles of 30 s on/30 s off and were centrifuged at 17,000 × g for 30 min at 4 °C. The supernatants were retained for protein measurement.

Nuclear and cytosolic fractions were prepared using an adapted fractionation protocol [36]. Tissue was homogenized on ice with a glass Dounce homogenizer in STM buffer containing 250 mM sucrose, 50 mM Tris-HCl (pH 7.4), 5 mM MgCl and Halt Protease and Phosphatase Inhibitor Cocktail. Following centrifugation at 800 × g for 15 min, the pellet was retained for nuclear extraction and the supernatant for cytosolic extraction. The cytosolic preparation was centrifuged at 800 × g for 10 min and then at 11,000 × g for 10 min. The supernatant was mixed 1:1 with 100% acetone, incubated at −20 °C for 2 h and centrifuged at 12,000 × g for 5 min. The pellet was resuspended in the STM buffer.

The nuclear preparation was resuspended in STM buffer and centrifuged at 500 × g for 15 min; the pellet was retained, resuspended in STM buffer, and centrifuged again at 1,000 × g for 15 min, with the pellet again retained for downstream nuclear extraction. The nuclear pellet was resuspended in NET buffer containing 20 mM HEPES (pH 7.9), 1.5 mM MgCl, 0.5 M NaCl, 0.2 mM EDTA, 20% glycerol, 1% Triton X-100 and Halt Protease and Phosphatase Inhibitor Cocktail. Samples were kept on ice for 30 min, sonicated three times for 10 s with 30 s intervals on ice, and centrifuged at 9,000 × g for 30 min. The final supernatant was retained as the nuclear extract. All centrifugation steps were performed at 4°C. Protein concentration was determined using the Pierce BCA Protein Assay Kit (Thermo Fisher Scientific, USA).

### Western blotting

Whole LZ lysates (10 µg per lane) and undiluted nuclear and cytosolic fractions (10 µL per lane) were separated on 10% gels using MOPS and MES running buffers, respectively, and transferred to 0.2 µm nitrocellulose membranes. Signals were acquired with an Odyssey CLx system and quantified using Image Studio 6.1.

KEAP1 (∼69 kDa) was normalized to β-actin, and nuclear and cytosolic NRF2 (∼95–100 kDa) to histone H2A and GAPDH, respectively. NRF2 localization was assessed using N/(N + C) × 100, where N and C represent the respective normalized signals. The detailed protocol, antibody information and full-length blots are provided in the Supplementary Methods, Supplementary Table S1 and Supplementary Figure S2, respectively. Measurements affected by persistent technical errors were excluded.

### Statistical analysis

RT-qPCR and Western blot data were analyzed using GraphPad Prism version 10.6.1. Group comparisons used unpaired t tests with Welch’s correction. Normality was assessed with Shapiro–Wilk tests, and the ROUT procedure with Q = 1% was used to identify outliers. RT-qPCR expression was first assessed with sexes pooled; when the pooled comparison was not significant, comparisons were performed within each sex. Data are presented as mean ± SEM, and P < 0.05 was considered statistically significant. NanoString comparisons used the FDR and fold-change criteria described above.

## Results

### Prenatal THC alters expression of metabolic and regulatory genes in the placental labyrinth

Targeted NanoString profiling identified a transcriptional response to prenatal THC exposure in the LZ at GD17.5. *Gls*, encoding glutaminase, *Keap1*, encoding Kelch-like ECH-associated protein 1, and *Flt3*, encoding FMS-like tyrosine kinase 3, were downregulated relative to VEH, with log_2_ fold changes of −0.259, −0.222 and −0.412, respectively. All three genes met the differential-expression criteria of FDR ≤0.10 and |log_2_ fold change| ≥0.2, with an FDR of 0.0904 for each gene (Fig. 1B). No genes met these criteria in the CBD-versus-VEH comparison (Fig. 1C). *Gls*, *Keap1* and *Flt3* were selected for RT-qPCR follow-up.

Fetal sex was included as a covariate in the NanoString differential-expression analysis. Follow-up RT-qPCR analyses of the selected targets examined expression in both sexes combined and separately in male and female placentas.

### Pathway analysis identifies transcriptional changes in arginine metabolism

Pathway-level analysis identified transcriptional changes in arginine metabolism following prenatal THC exposure in the placental LZ. Among the 34 metabolic pathways tested, this nine-gene set met the significance criterion in the THC-versus-VEH comparison (Mixed-test P = 0.002; FDR = 0.0527; Fig. 2A). Within this set, the expression pattern included lower *Gls* and *Gls2* expression alongside higher *Got1* and *Got2* expression (Fig. 2B). *Gls*, *Got2*, *Gls2*, *Got1* and *Nos2* made the largest contributions to the pathway-level statistic. The arginine-metabolism finding was retained when the analysis was extended to all 36 annotated categories (FDR = 0.0558). No pathway met the FDR ≤0.10 criterion in the CBD-versus-VEH comparison.

**Figure 2.**
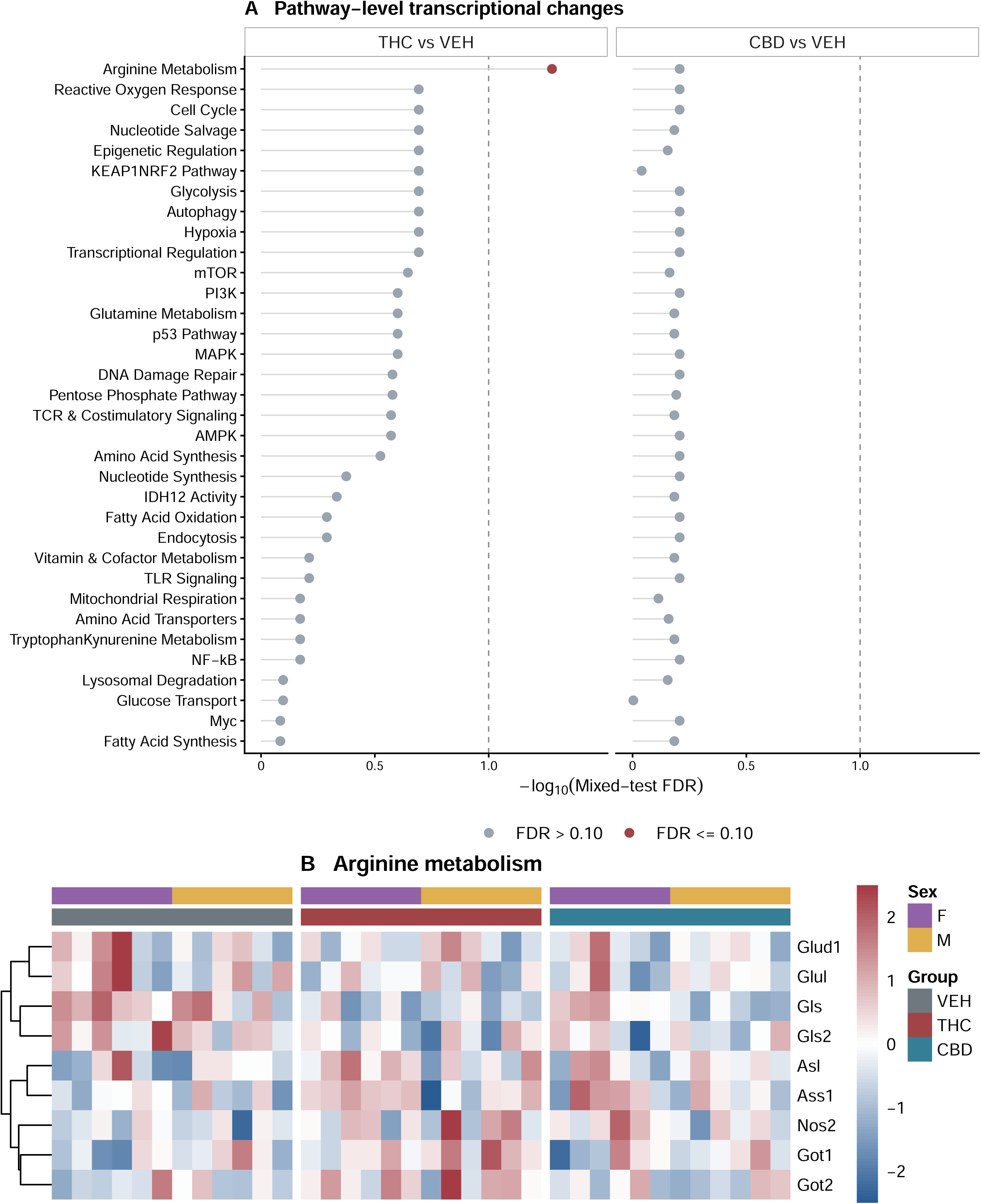
Pathway analysis identifies transcriptional changes in arginine metabolism. (A) −log_10_ Mixed-test FDR values for 34 metabolic pathways in sex-adjusted THC-versus-VEH and CBD-versus-VEH comparisons using the 463-gene matrix. The dashed line indicates FDR = 0.10; red points meet this criterion. (B) Heatmap of all nine tested genes annotated to arginine metabolism across 36 placentas. Colors show gene-wise z-scores of normalized log_2_ expression, capped at ±2.5 for display. Columns represent individual placentas ordered by treatment and fetal sex; rows are hierarchically clustered. Each group contains six female-associated and six male-associated placentas.

### Transcript measurements confirm Keap1 downregulation and identify Gls and Flt3 reductions in male samples

RT-qPCR confirmed lower *Keap1* expression in THC-exposed LZ samples with sexes pooled (t(22.00) = 2.84, P = 0.010; n = 12 placentas per group; Fig. 3A). This agreement between NanoString and RT-qPCR identifies *Keap1* as a consistent transcriptional target in the THC-exposed placenta. Follow-up measurements focused on THC and VEH, based on the expression pattern identified by the NanoString screen.

**Figure 3.**
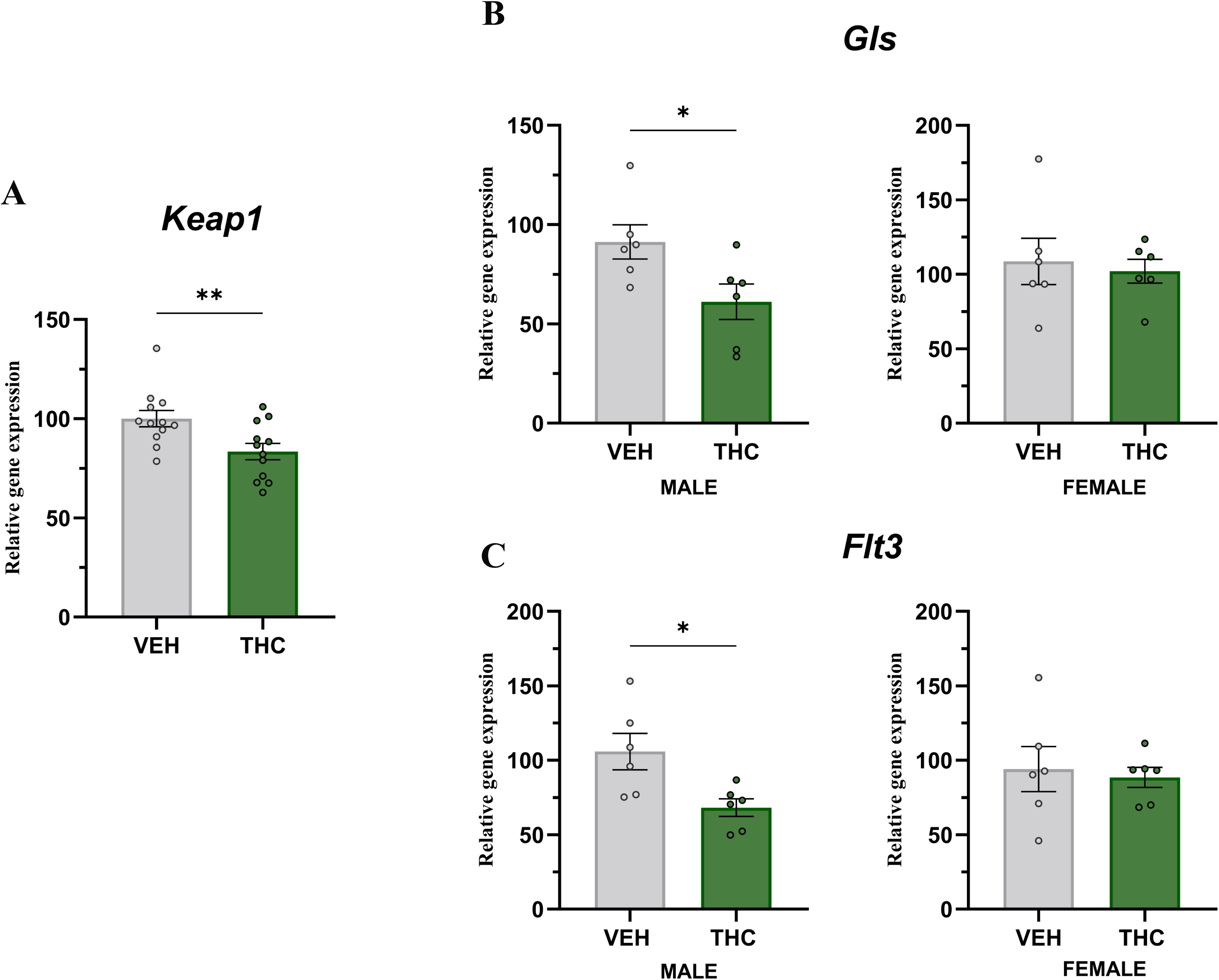
Quantitative PCR analysis of selected placental transcripts. Relative expression of (A) *Keap1* with sexes pooled, (B) *Gls* stratified by fetal sex, and (C) *Flt3* stratified by fetal sex at GD17.5. Points represent individual placentas. Values are mean ± SEM. Sample sizes were 12 placentas per group for *Keap1* and six per group per sex for *Gls* and *Flt3*. Statistical comparisons used two-sided unpaired t tests with Welch’s correction. *P < 0.05; **P < 0.01.

With sexes pooled, *Gls* (t(21.94) = 1.50, P = 0.147) and *Flt3* (t(17.18) = 2.01, P = 0.061) expression did not differ significantly between THC and VEH groups (n = 12 placentas per group). Sex-stratified comparisons revealed lower *Gls* expression (t(9.99) = 2.43, P = 0.035) and lower *Flt3* expression (t(7.19) = 2.78, P = 0.027) in male-associated placentas after THC exposure (n = 6 per group; Fig. 3B,C). The corresponding comparisons in female-associated placentas were not significant (both P > 0.7).

### KEAP1 protein increases without detectable NRF2 redistribution or Hmox1 induction

The reduction in *Keap1* transcript abundance prompted examination of KEAP1 protein and downstream NRF2-related readouts. KEAP1 regulates NRF2, a transcription factor that controls antioxidant-response genes including *Hmox1* [37,38]. KEAP1 was quantified with sexes pooled, whereas nuclear and cytosolic NRF2 fractions were analyzed separately by sex owing to gel-loading capacity. *Hmox1* expression was assessed separately by sex, in parallel with the NRF2 fractions.

KEAP1 protein abundance was higher in THC-exposed placentas than in VEH controls (mean difference = 0.0067 AU, 95% CI [0.0032, 0.0101]; Welch’s t(16.51) = 4.09, P < 0.001, η² = 0.50; n = 12 VEH, 11 THC placentas; Fig. 4A,B). The increase in protein abundance contrasted with the reduction in *Keap1* mRNA, revealing divergent transcript and protein responses to prenatal THC exposure.

**Figure 4.**
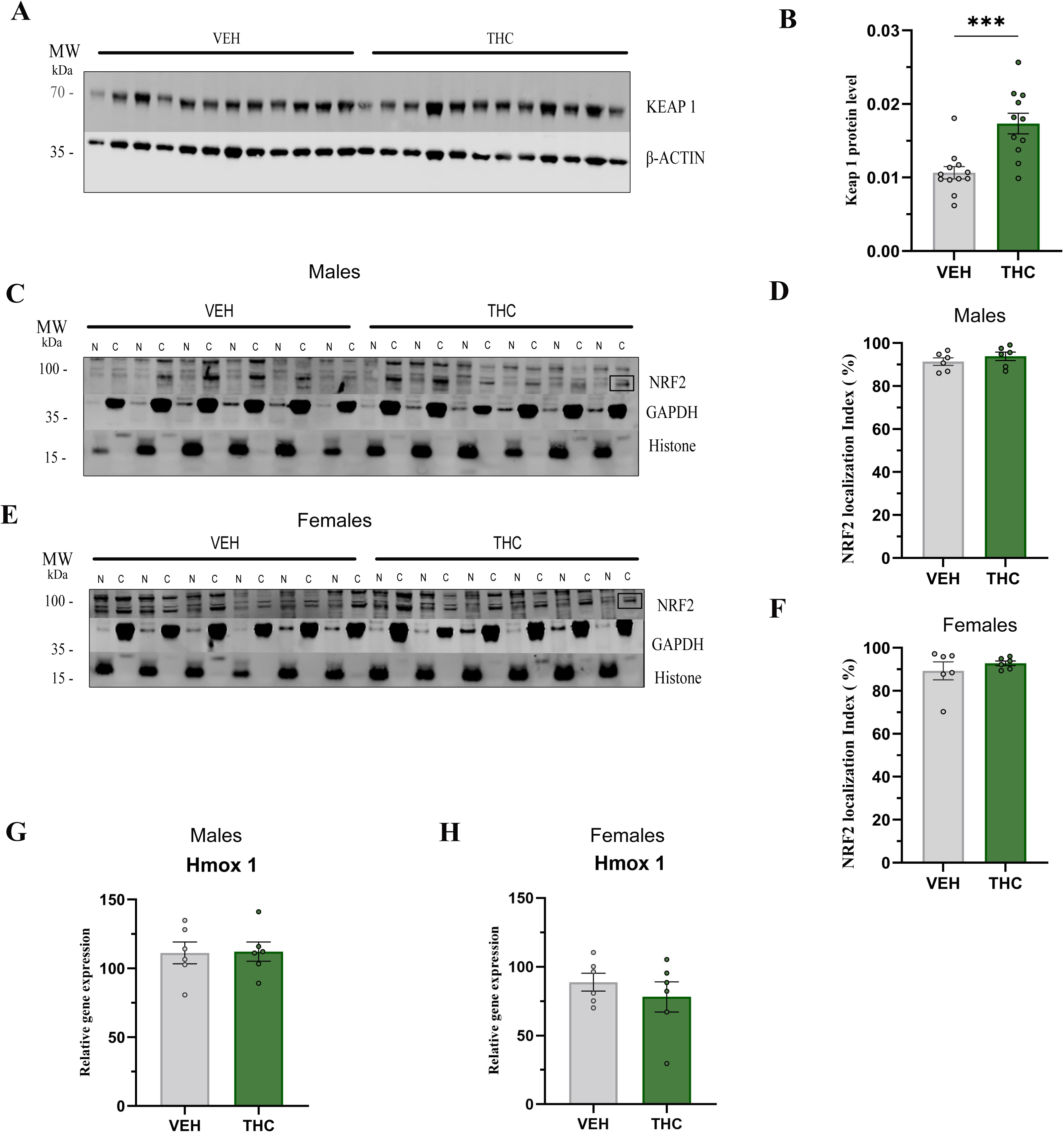
KEAP1 abundance and NRF2 measurements after prenatal THC exposure. (A) Representative KEAP1 and β-actin immunoblots from LZ lysates. (B) KEAP1 abundance normalized to β-actin (n = 12 VEH and 11 THC placentas; one THC sample was excluded owing to poor blot quality). (C,E) Representative nuclear (N) and cytosolic (C) extracts from male– and female-associated placentas, respectively, probed for NRF2, histone H2A and GAPDH. (D,F) NRF2 localization index in males and females, calculated as N/(N + C) × 100. (G,H) *Hmox1* expression in males and females. For NRF2 and *Hmox1*, n = 6 placentas per group per sex. Values are mean ± SEM. ***P < 0.001 for KEAP1, using an unpaired t test with Welch’s correction. Full-length blots are shown in Supplementary Figure S2.

The NRF2 localization index showed no significant difference between THC and VEH in males (mean difference = 2.50%, 95% CI [−3.38, 8.38]; Welch’s t(9.87) = 0.95, P = 0.366, η² = 0.08) or females (mean difference = 3.46%, 95% CI [−7.23, 14.14]; Welch’s t(5.67) = 0.80, P = 0.455, η² = 0.10; n = 6 placentas per group per sex; Fig. 4C–F). Nuclear and cytosolic NRF2 levels analyzed separately also showed no significant differences.

*Hmox1* expression followed the same pattern, with no significant difference in males (mean difference = 0.92, 95% CI [−22.62, 24.47]; Welch’s t(9.84) = 0.09, P = 0.932, η² < 0.01) or females (mean difference = −10.79, 95% CI [−40.16, 18.58]; Welch’s t(8.08) = 0.85, P = 0.422, η² = 0.08; n = 6 placentas per group per sex; Fig. 4G,H). Thus, the KEAP1 protein increase was not accompanied by a detectable change in the assessed NRF2 or *Hmox1* readouts at GD17.5.

## Discussion

Prenatal THC exposure altered amino acid metabolic gene regulation in the placental LZ at GD17.5. NanoString profiling identified lower *Gls*, *Keap1* and *Flt3* expression and a transcriptional response within arginine metabolism. RT-qPCR confirmed lower *Keap1* expression and identified *Gls* and *Flt3* reductions in male samples. KEAP1 protein abundance increased despite lower *Keap1* mRNA, while NRF2 localization and *Hmox1* expression showed no significant differences. These findings connect an amino acid metabolic expression response with regulation of KEAP1 in the placental exchange region.

The LZ contains the trophoblast and vascular structures responsible for maternal–fetal exchange, making its metabolic regulation directly relevant to the fetal environment [23,28]. Previous studies identified labyrinth vascular abnormalities after THC or CBD exposure [14,18], and layer-enriched analyses reveal responses obscured in whole placental tissue [29]. Our findings extend this work to a defined metabolic gene set and KEAP1 protein regulation.

The arginine-metabolism result places *Gls* within a broader amino acid expression response. The nine-gene set encompasses glutamine and glutamate handling alongside arginine-related processes. Lower *Gls* and *Gls2* expression occurred together with higher *Got1* and *Got2* expression, indicating changes across several components of the set. This pathway-level view connects individual transcript targets with a wider metabolic response in the placental exchange region.

*Gls* links the pathway result with the transcript measurements. It encodes glutaminase, which converts glutamine to glutamate; glutamine contributes to mitochondrial respiration in primary human trophoblasts [25]. The placenta also mediates glutamine–glutamate exchange between maternal and fetal circulations [39], linking glutaminase to trophoblast energy metabolism and fetal amino acid supply. Lower *Gls* expression in NanoString and male RT-qPCR samples complements evidence connecting THC with altered placental nutrient exchange and trophoblast function [14,40].

The THC group showed gene-level and arginine-metabolism responses, whereas no genes or pathways met the corresponding criteria in the CBD group. Their different receptor engagement and signaling provide context for this screening pattern [8,9]. Gestational CBD affects vascular development, trophoblast metabolism and transport [18–20,22], emphasizing that cannabinoid responses depend on exposure conditions and the placental processes examined. Here, the findings direct attention to THC-associated metabolic and stress-response gene regulation.

THC affects trophoblast turnover and differentiation, induces endoplasmic reticulum stress and alters mitochondrial function [16,17,40,41]. STAT3 signaling and trophoblast differentiation also respond to THC or cannabis smoke extract [15,42]. These processes are linked to the energetic and redox demands of placental development, providing a biological context for the amino acid expression pattern and *Keap1* response.

The opposite directions of *Keap1* mRNA and KEAP1 protein point to differential regulation at transcript and protein levels, potentially involving translation or protein stability. KEAP1 controls NRF2 and participates in stress-response processes beyond canonical NRF2 signaling [43]. Increased KEAP1 abundance may therefore represent an adaptive component of repeated THC exposure. Cannabinoid effects on p62/SQSTM1-associated pathways and CBD-induced changes in trophoblast autophagy provide further context for these divergent transcript and protein responses [44–46].

The KEAP1 increase occurred without a detectable NRF2 localization shift or *Hmox1* induction at GD17.5, placing KEAP1 abundance at the center of the measured protein response. Cannabinoid regulation of NRF2 varies across experimental systems [47]; THC and CBD, for example, produce different NRF2-related transcriptional profiles in microglial cells [48]. The combined measurements identify KEAP1 abundance as a feature of the placental response to gestational THC.

*Flt3* adds a developmental dimension to this response. It is associated with hematopoietic progenitor biology [49], and the placenta supports hematopoietic development [50]. Its downregulation in NanoString and male RT-qPCR samples raises the possibility that prenatal THC influences gene regulation in cell populations contributing to this role, alongside its amino acid metabolic and KEAP1-related responses.

The *Gls* and *Flt3* reductions in male samples place fetal sex within the observed placental response. Placental physiology varies with sex and developmental context [51,52], with sex-related patterns reported after gestational hypoxia, maternal obesity [53,54] and THC exposure [55]. Our findings add selected expression changes within the placental exchange region to this literature.

Integrating pathway analysis with transcript and protein measurements links prenatal THC exposure to molecular responses in a region central to maternal–fetal communication. Amino acid metabolic gene expression and KEAP1 regulation provide targets for investigating placental adaptation to cannabinoids and its relevance to the placenta–brain axis [5,6].

### Limitations

The study examined one intraperitoneal dose and one gestational time point in LZ-enriched tissue using a targeted expression panel. The small number of litters and multiple placentas sampled per dam limit statistical independence and precision, particularly in sex-stratified comparisons. *Hmox1* was the only downstream NRF2 target measured, and mitochondrial NRF2 was not assessed.

### Conclusion

Prenatal THC exposure altered arginine-metabolism gene expression and reduced *Gls*, *Keap1* and *Flt3* transcripts in the mouse placental labyrinth. Follow-up measurements confirmed lower *Keap1* expression, identified *Gls* and *Flt3* reductions in male samples, and showed increased KEAP1 protein abundance. These findings identify amino acid metabolic gene regulation and KEAP1 regulation as complementary features of placental responses to prenatal THC.

## CRediT authorship contribution statement

**Eleni Sia:** Investigation, Formal analysis, Visualization, Writing – original draft.

**Pedro Henrique Gobira:** Investigation, Writing – review & editing.

**Fenghua Chen:** Investigation (animal experiments), Writing – review & editing.

**Gülendam Sezer:** Investigation (molecular experiments).

**Heidi Kaastrup Müller:** Investigation (protein assays).

**Sâmia Joca:** Conceptualization, Supervision.

**Cristiana Cruceanu:** Conceptualization, Supervision.

**Rodrigo Grassi-Oliveira:** Conceptualization, Methodology, Investigation, Formal analysis, Visualization, Writing – original draft, Writing – review & editing, Supervision, Project administration, Funding acquisition.

## Data availability

NanoString raw and normalized expression data, sample metadata, and complete gene– and pathway-level results are provided in Supplementary Data S1 and S2. Full-length Western blot images are provided in Supplementary Figure S2.

## Declaration of generative AI and AI-assisted technologies in the manuscript preparation process

ChatGPT (OpenAI) was used to assist with manuscript editing and organization and with the preparation of analysis scripts. The authors reviewed and edited the content as needed and take full responsibility for the content of the published article.

## References

[1] K.C. Young-Wolff, et al. Rates of prenatal cannabis use among pregnant women before and during the COVID-19 pandemic. JAMA 326 (2021) 1745–1747.

[2] N.D. Volkow, B. Han, W.M. Compton, E.F. McCance-Katz, Self-reported medical and nonmedical cannabis use among pregnant women in the United States. JAMA 322 (2019) 167– 169.

[3] C.D. Azubuike, O. Grundmann, A.J. Goodin, Neurodevelopmental effects of perinatal exposure to cannabis on progeny: A narrative review. Drug Alcohol Depend. Rep. 16 (2025) 100372.

[4] P. Díaz, T.L. Powell, T. Jansson, The role of placental nutrient sensing in maternal-fetal resource allocation. Biol. Reprod. 91 (2014) 82.

[5] C.S. Rosenfeld, The placenta brain axis. J. Neurosci. Res. 99 (2021) 271–283.

[6] H. Hazzard, B. Blaine, B.B. Borrman, S.Q. Shen, S.B. Gumusoglu, Extracellular vesicles and the placenta-brain axis: Mechanisms of shared risk in pregnancy and perinatal psychiatry. Curr. Opin. Physiol. 47 (2026) 100913.

[7] J. Maia, B.M. Fonseca, N. Teixeira, G. Correia-da-Silva, The fundamental role of the endocannabinoid system in endometrium and placenta: implications in pathophysiological aspects of uterine and pregnancy disorders. Hum. Reprod. Update 26 (2020) 586–602.

[8] A.C.E. Rokeby, B.V. Natale, D.R.C. Natale, Cannabinoids and the placenta: Receptors, signaling and outcomes. Placenta 135 (2023) 51–61.

[9] N. Stella, THC and CBD: Similarities and differences between siblings. Neuron 111 (2023) 302– 327.

[10] M.S. Harhangi, et al. Recreational cannabis use during human pregnancy: Its effects on the placenta and endocannabinoid system. Int. J. Mol. Sci. 27 (2026) 1398.

[11] J. Maia, et al. Effects of cannabis tetrahydrocannabinol on endocannabinoid homeostasis in human placenta. Arch. Toxicol. 93 (2019) 649–658.

[12] T. Podinic, et al. Prenatal cannabis smoke exposure alters placental development in a murine model of pregnancy. PLoS One 21 (2026) e0328123.

[13] V.H.J. Roberts, et al. Chronic prenatal delta-9-tetrahydrocannabinol exposure adversely impacts placental function and development in a rhesus macaque model. Sci. Rep. 12 (2022) 20260.

[14] B.V. Natale, et al. Δ^9^-tetrahydrocannabinol exposure during rat pregnancy leads to symmetrical fetal growth restriction and labyrinth-specific vascular defects in the placenta. Sci. Rep. 10 (2020) 544.

[15] X. Chang, et al. Suppression of STAT3 signaling by Δ^9^-tetrahydrocannabinol (THC) induces trophoblast dysfunction. Cell. Physiol. Biochem. 42 (2017) 537–550.

[16] O.S. Walker, et al. Delta-9-tetrahydrocannabinol inhibits invasion of HTR8/SVneo human extravillous trophoblast cells and negatively impacts mitochondrial function. Sci. Rep. 11 (2021) 4029.

[17] O.S. Walker, et al. Delta-9-tetrahydrocannabinol disrupts mitochondrial function and attenuates syncytialization in human placental BeWo cells. Physiol. Rep. 8 (2020) e14476.

[18] S. Allen, et al. Cannabidiol exposure during rat pregnancy leads to labyrinth-specific vascular defects in the placenta and reduced fetal growth. Cannabis Cannabinoid Res. 9 (2024) 766–780.

[19] T.M. Ritchie, et al. The impact of oral cannabis consumption during pregnancy on maternal spiral artery remodelling, fetal growth and offspring behaviour in mice. EBioMedicine 114 (2025) 105572.

[20] T. Podinic, et al. Cannabidiol disrupts mitochondrial respiration and metabolism and dysregulates trophoblast cell differentiation. Cells 13 (2024) 486.

[21] R. Portillo, et al. Cannabidiol disrupts tryptophan metabolism in the human term placenta. Toxicology 505 (2024) 153813.

[22] V. Feinshtein, et al. Cannabidiol enhances xenobiotic permeability through the human placental barrier by direct inhibition of breast cancer resistance protein: an ex vivo study. Am. J. Obstet. Gynecol. 209 (2013) 573.e1–573.e15.

[23] I.L.M.H. Aye, C.E. Aiken, D.S. Charnock-Jones, G.C.S. Smith, Placental energy metabolism in health and disease-significance of development and implications for preeclampsia. Am. J. Obstet. Gynecol. 226 (2022) S928–S944.

[24] X. Yu, et al. Acetyl-CoA metabolism maintains histone acetylation for syncytialization of human placental trophoblast stem cells. Cell Stem Cell 31 (2024) 1280–1297.e7.

[25] Y. Wang, M. Bucher, L. Myatt, Use of glucose, glutamine, and fatty acids for trophoblast respiration in lean women, women with obesity, and women with gestational diabetes. J. Clin. Endocrinol. Metab. 104 (2019) 4178–4187.

[26] A. Busquets-García, J.P. Bolaños, G. Marsicano, Metabolic messengers: Endocannabinoids. Nat. Metab. 4 (2022) 848–855.

[27] A.N. Sferruzzi-Perri, J. Lopez-Tello, E. Salazar-Petres, Placental adaptations supporting fetal growth during normal and adverse gestational environments. Exp. Physiol. 108 (2023) 371–397.

[28] S.A. Elmore, et al. Histology atlas of the developing mouse placenta. Toxicol. Pathol. 50 (2022) 60–117.

[29] J.C. Hercus, et al. The whole is lesser than the sum of its parts? Dissecting layer-enriched samples of rodent placenta is worth the effort. Placenta 157 (2024) 76–80.

[30] T. Black, et al. Characterization of cannabinoid plasma concentration, maternal health, and cytokine levels in a rat model of prenatal Cannabis smoke exposure. Sci. Rep. 13 (2023) 21070.

[31] R. Behringer, M. Gertsenstein, K.V. Nagy, A. Nagy, Preparation of polymerase chain reaction template DNA from mouse tail tissue. Cold Spring Harb. Protoc. 2019 (2019) db.prot092700.

[32] D. Qu, A. McDonald, K.J. Whiteley, S.A. Bainbridge, S.L. Adamson, Layer-enriched tissue dissection of the mouse placenta in late gestation, in: The Guide to Investigation of Mouse Pregnancy, Elsevier, 2014, pp. 529–536.

[33] D. Wu, et al. ROAST: rotation gene set tests for complex microarray experiments. Bioinformatics 26 (2010) 2176–2182.

[34] M.E. Ritchie, et al. limma powers differential expression analyses for RNA-sequencing and microarray studies. Nucleic Acids Res. 43 (2015) e47.

[35] B. Elfving, Investigation of synaptic vesicle proteins in rat brain tissue using real-time qPCR. Methods Mol. Biol. 2417 (2022) 59–68.

[36] I. Dimauro, T. Pearson, D. Caporossi, M.J. Jackson, A simple protocol for the subcellular fractionation of skeletal muscle cells and tissue. BMC Res. Notes 5 (2012) 513.

[37] G. Tossetta, et al. Modulation of NRF2/KEAP1 signaling in preeclampsia. Cells 12 (2023) 1545.

[38] O.A.G. Tantengco, et al. The role of nuclear factor erythroid 2-related factor 2 (NRF2) in normal and pathological pregnancy: A systematic review. Am. J. Reprod. Immunol. 86 (2021) e13496.

[39] X. Wu, et al. Glutamate–glutamine cycle and exchange in the placenta–fetus unit during late pregnancy. Amino Acids 47 (2015) 45–53.

[40] M.A. Costa, B.M. Fonseca, F. Marques, N.A. Teixeira, G. Correia-da-Silva, The psychoactive compound of Cannabis sativa, Δ(9)-tetrahydrocannabinol (THC) inhibits the human trophoblast cell turnover. Toxicology 334 (2015) 94–103.

[41] T. Lojpur, et al. Δ^9^-Tetrahydrocannabinol leads to endoplasmic reticulum stress and mitochondrial dysfunction in human BeWo trophoblasts. Reprod. Toxicol. 87 (2019) 21–31.

[42] C. Monaco, et al. Cannabis smoke extract disrupts trophoblast differentiation and causes mitochondrial dysfunction beyond the effects of Δ^9^-THC alone. Sci. Rep. 16 (2026) 6253.

[43] A.T. Dinkova-Kostova, I.M. Copple, Advances and challenges in therapeutic targeting of NRF2. Trends Pharmacol. Sci. 44 (2023) 137–149.

[44] C. Blázquez, et al. Cannabinoid-induced motor dysfunction via autophagy inhibition. Autophagy 16 (2020) 2289–2291.

[45] F. Armeli, B. Mengoni, D.L. Laskin, R. Businaro, Interplay among oxidative stress, autophagy, and the endocannabinoid system in neurodegenerative diseases: Role of the Nrf2-p62/SQSTM1 pathway and nutraceutical activation. Curr. Issues Mol. Biol. 46 (2024) 6868–6884.

[46] P. Alves, C. Amaral, N. Teixeira, G. Correia-da-Silva, Cannabidiol disrupts apoptosis, autophagy and invasion processes of placental trophoblasts. Arch. Toxicol. 95 (2021) 3393–3406.

[47] P. Marini, M. Maccarrone, L. Saso, P. Tucci, The effect of phytocannabinoids and endocannabinoids on Nrf2 activity in the central nervous system and periphery. Neurol. Int. 16 (2024) 776–789.

[48] A. Juknat, et al. Differential transcriptional profiles mediated by exposure to the cannabinoids cannabidiol and Δ^9^-tetrahydrocannabinol in BV-2 microglial cells. Br. J. Pharmacol. 165 (2012) 2512–2528.

[49] O. deLapeyrière, et al. Expression of *Flt3* tyrosine kinase receptor gene in mouse hematopoietic and nervous tissues. Differentiation 58 (1995) 351–359.

[50] C. Gekas, et al. Hematopoietic stem cell development in the placenta. Int. J. Dev. Biol. 54 (2010) 1089–1098.

[51] A.S. Meakin, J.S.M. Cuffe, J.R.T. Darby, J.L. Morrison, V.L. Clifton, Let’s talk about placental sex, baby: Understanding mechanisms that drive female– and male-specific fetal growth and developmental outcomes. Int. J. Mol. Sci. 22 (2021) 6386.

[52] C. Stenhouse, F.W. Bazer, C.J. Ashworth, Sexual dimorphism in placental development and function: Comparative physiology with an emphasis on the pig. Mol. Reprod. Dev. 90 (2023) 684–696.

[53] H. Song, B.P. Telugu, L.P. Thompson, Sexual dimorphism of mitochondrial function in the hypoxic guinea pig placenta. Biol. Reprod. 100 (2019) 208–216.

[54] T. Napso, et al. Diet-induced maternal obesity impacts feto-placental growth and induces sex-specific alterations in placental morphology, mitochondrial bioenergetics, dynamics, lipid metabolism and oxidative stress in mice. Acta Physiol. (Oxf.) 234 (2022) e13795.

[55] A.M. Kocsis, et al. Identifying established human placental markers of schizophrenia in rodents after gestational Δ9-tetrahydrocannabinol exposure. Biol. Reprod. 114 (2026) 246–258.

