## Supplementary for "Prenatal THC exposure alters metabolic gene expression and KEAP1 regulation in the mouse placental labyrinth": Supplementary_Information.pdf

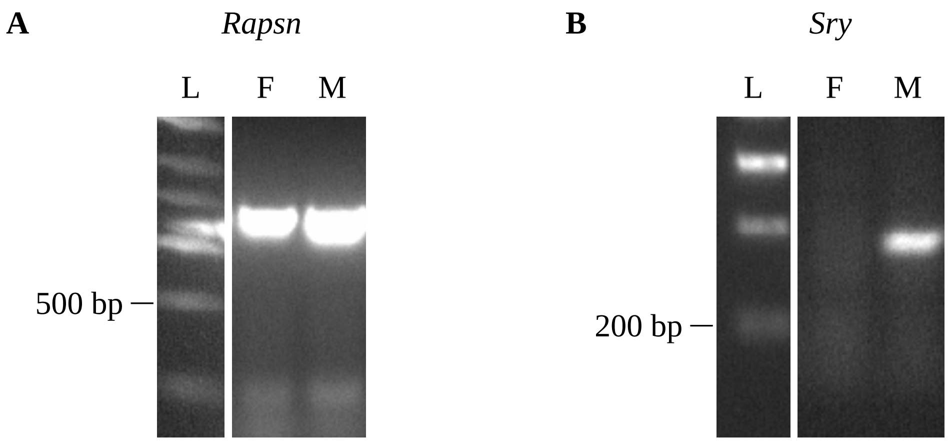


**Supplementary Figure S1.** PCR-based sex determination of mouse fetuses using tail-derived DNA collected at GD17.5. Representative female (F) and male (M) samples are shown. (A) Autosomal *Rapsn* (590 bp), amplified in both sexes as a DNA control. (B) Y chromosome-specific *Sry* (230 bp), amplified only in the male sample. The same DNA samples were used for both reactions. Sex was determined for all fetuses (n = 6 per sex per group). L, DNA ladder.

### Supplementary Methods

#### Western blotting

Whole placental labyrinth zone (LZ) lysates were adjusted to 1 µg/µL, and 10 µg of protein (10 µL) was loaded per lane. Nuclear and cytosolic fractions were not diluted, and equal volumes (10 µL) were loaded per lane.

All samples were mixed with sample buffer containing DTT, incubated at 65 °C for 20 min and separated on 10% 26-well gels using MOPS running buffer for whole lysates or MES running buffer for subcellular fractions. Proteins were transferred to 0.2 µm nitrocellulose membranes using a Trans-Blot Turbo system (Bio-Rad, USA) for 7 min. PageRuler Plus prestained markers were used to estimate molecular weights.

Membranes were blocked for 1 h with Intercept TBS Blocking Buffer and incubated with primary antibodies overnight at 4 °C. After washing in TBS containing 0.1% Tween 20, membranes were incubated with the appropriate secondary antibodies (LI-COR Biosciences, USA) for 1 h. Primary antibodies and dilutions, and secondary antibody identities, are listed in Supplementary Table S1.

Signals were acquired with an Odyssey CLx system and quantified using Image Studio software version 6.1. Bands were quantified at approximately 69 kDa (KEAP1), 42 kDa (β-actin), 95-100 kDa (NRF2), 37 kDa (GAPDH) and 14 kDa (histone H2A). KEAP1 was normalized to β-actin. Nuclear and cytosolic NRF2 signals were normalized to histone H2A and GAPDH, respectively. NRF2 localization was assessed using $N/(N + C) \times 100$, where N and C represent the respective normalized signals. Measurements affected by persistent technical errors were excluded. Full-length blots corresponding to Fig. 4 are shown in Supplementary Figure S2.

**Supplementary Table S1.** Antibodies used for western blotting (WB) of mouse placental tissue.

| Target | Catalog no. | Supplier | Host species | Dilution |
|---|---|---|---|---|
| *Housekeeping proteins* | | | | |
| Histone H2A | 2718 | Cell Signaling Technology | Rabbit | 1:500 |
| GAPDH | 42216 | LI-COR Biosciences | Rabbit | 1:500 |
| β-actin | 42212 | LI-COR Biosciences | Mouse | 1:3000 |
| *Target proteins* | | | | |
| KEAP1 | 8047T | Cell Signaling Technology | Rabbit | 1:1000 |
| NRF2 | 61812 | Cell Signaling Technology | Rabbit | 1:500 |

Secondary antibodies: IRDye® 800CW goat anti-rabbit IgG and IRDye® 680RD goat anti-mouse IgG (LI-COR Biosciences, USA).

**Supplementary Table S2.** Primer sequences used for RT-qPCR and sex-determination PCR.

| Gene | Gene name | Accession no.[a] | Forward primer (5′–3′) | Reverse primer (5′–3′) |
|---|---|---|---|---|
| *Housekeeping genes* | | | | |
| *Gapdh* | Glyceraldehyde-3-phosphate dehydrogenase | NM_008084.3 | TCACCACCATGGAGAAGGC | GCTAAGCAGTTGGTGGTGCA |
| *Rpl13a* | Ribosomal protein L13A | NM_009438.5 | ACAAGAAAAAGCGGATGGTG | TTCCGATAGTGCATCTTGGC |
| *Ywhaz* | Tyrosine 3-monooxygenase/ tryptophan 5-monooxygenase activation protein, zeta | BC089334 | TTGAGCAGAAGACGGAAGGT | GAAGCATTGGGGATCAAGAA |
| *Target genes* | | | | |
| *Gls* | Glutaminase | NM_001081081.2 | AACAGCGAGGGCAAGGAGAT | CAATCCTGTAGATTTGAGTGCTGTA |
| *Keap1* | Kelch-like ECH-associated protein 1 | NM_016679.4 | ATCTACGTCCTCGGAGGCTAT | TTGCTTCCGACAGGGTTCC |
| *Flt3* | FMS-like tyrosine kinase 3 | NM_010229.2 | TACAAACCAAGACCTGCCTGT | TCCTCGCACCATTCGGT |
| *Hmox1* | Heme oxygenase 1 | NM_010442.2 | GACAGCCCCACCAAGTTCAA | CAGCTCCTCAAACAGCTCAATG |
| *Sex-determination genes*[b] | | | | |
| *Sry* | Sex-determining region Y | – | TGACTGGGATGCAGTAGTTC | TGTGCTAGAGAGAAACCCTG |
| *Rapsn* | Receptor-associated protein of the synapse | – | AGGACTGGGTGGCTTCCAACTCCCAGACAC | AGCTTCTCATTGCTGCGCGCCAGGTTCAGG |

[a]Accession numbers refer to NCBI RefSeq/GenBank entries used for primer design.
[b]Genotyping PCR conditions: 95 °C for 5 min; 35 cycles of 95 °C for 30 s, 65 °C for 30 s and 72 °C for 45 s; final extension at 72 °C for 10 min.

**A** KEAP1, 800 nm channel

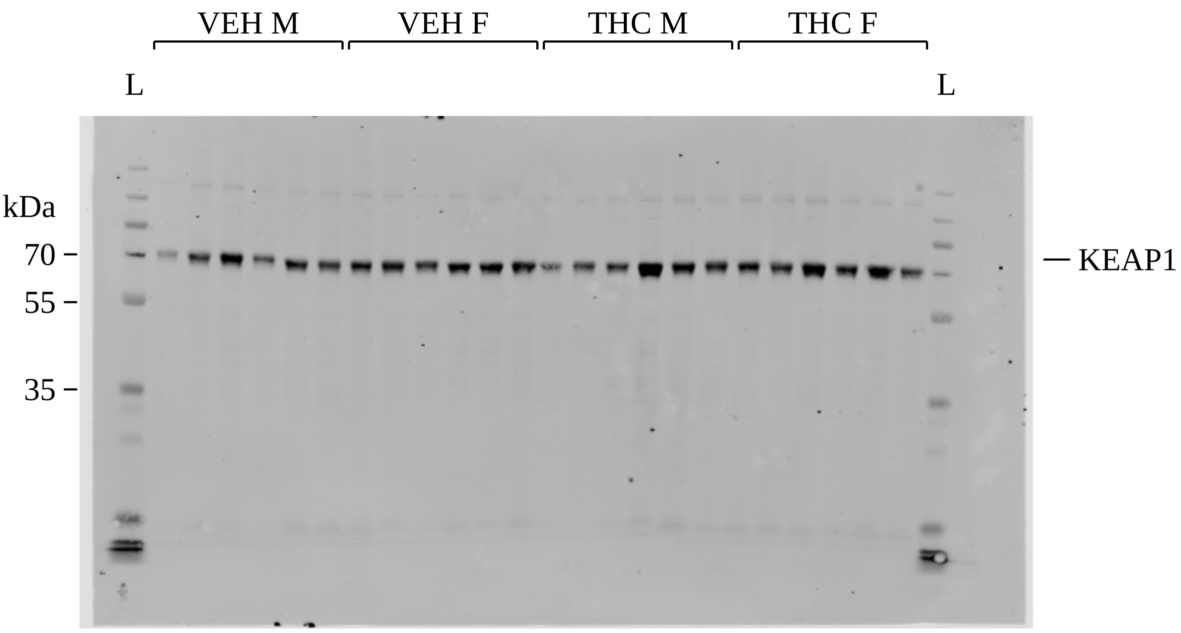


**B** β-actin, 700 nm channel

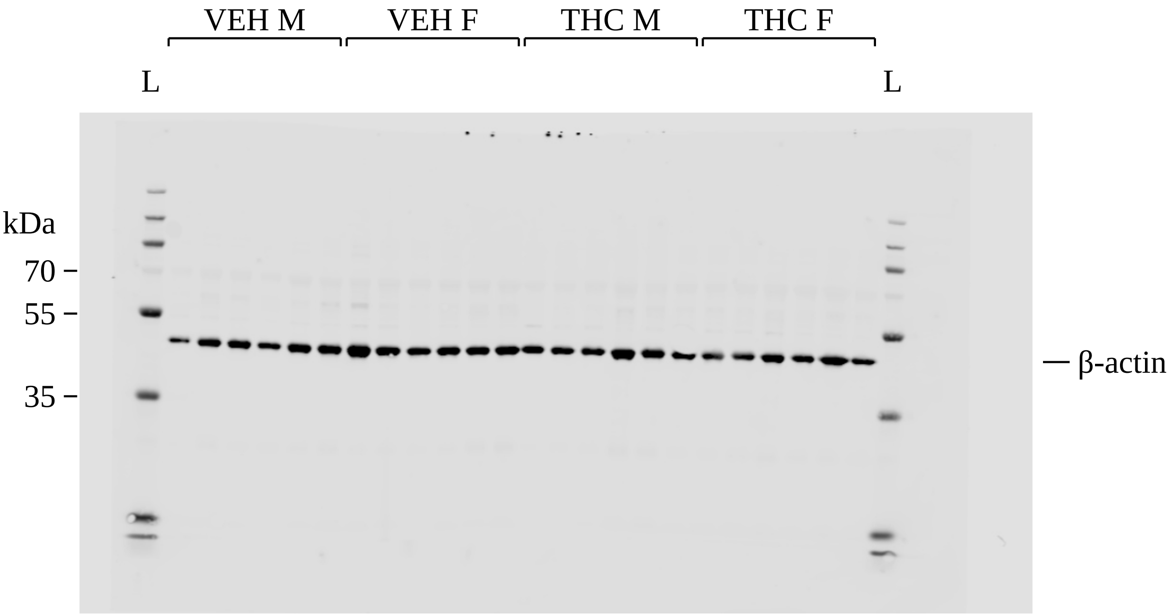

**C** NRF2 and GAPDH, 800 nm channel

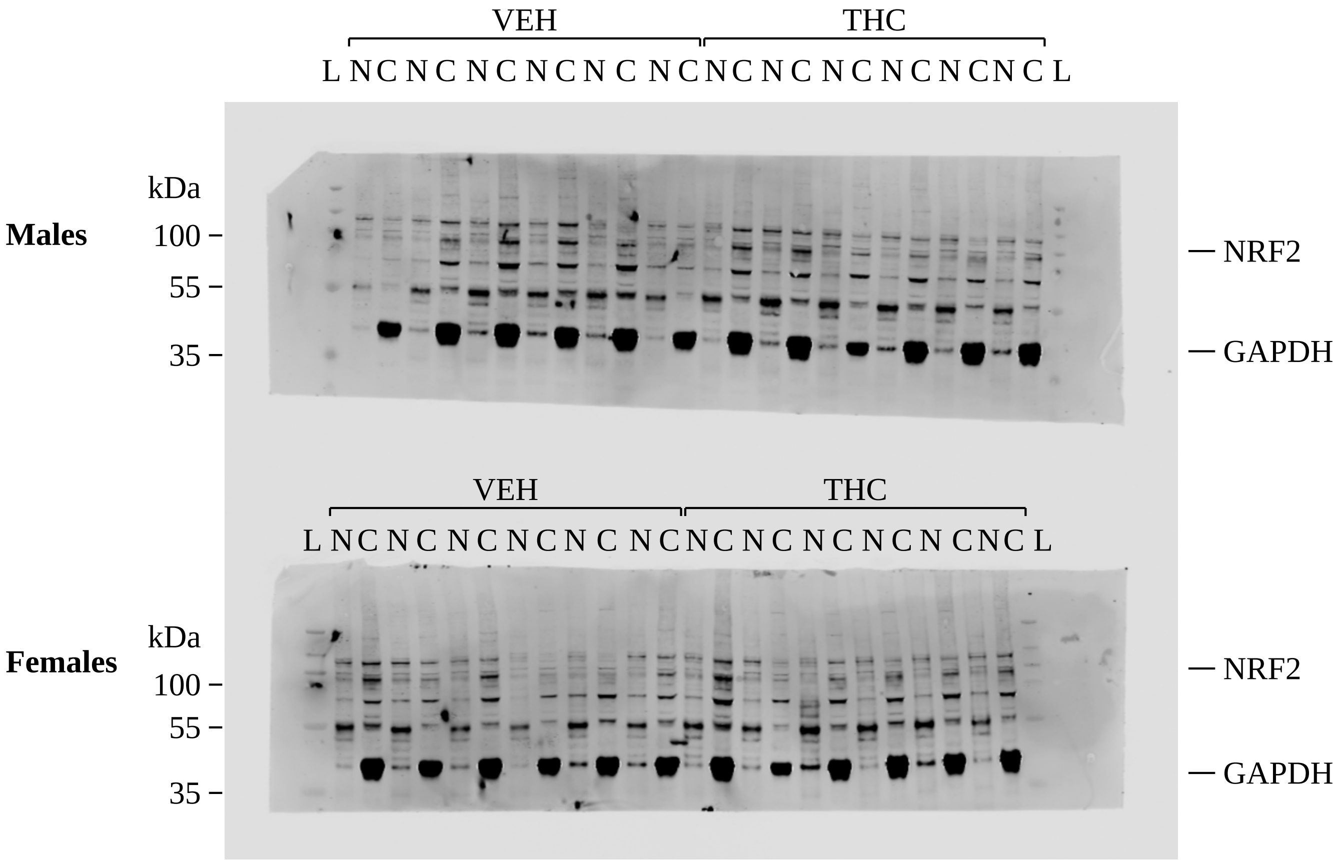

**D** Histone H2A, 800 nm channel

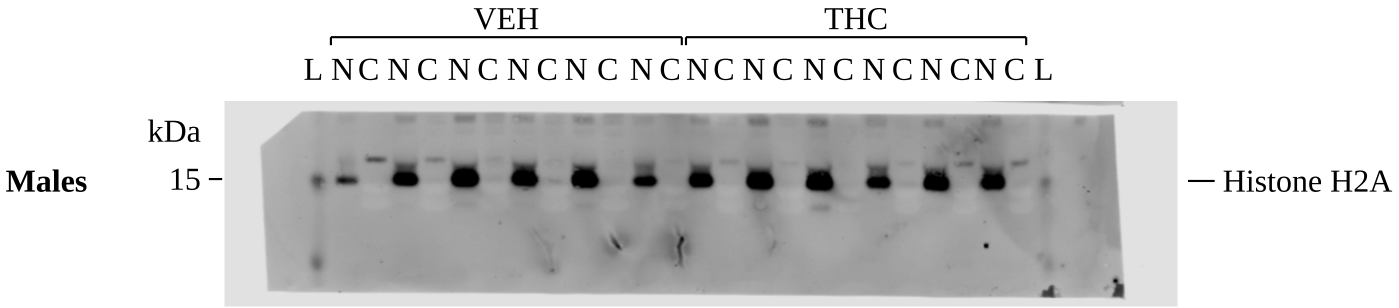


**E** Histone H2A, 800 nm channel

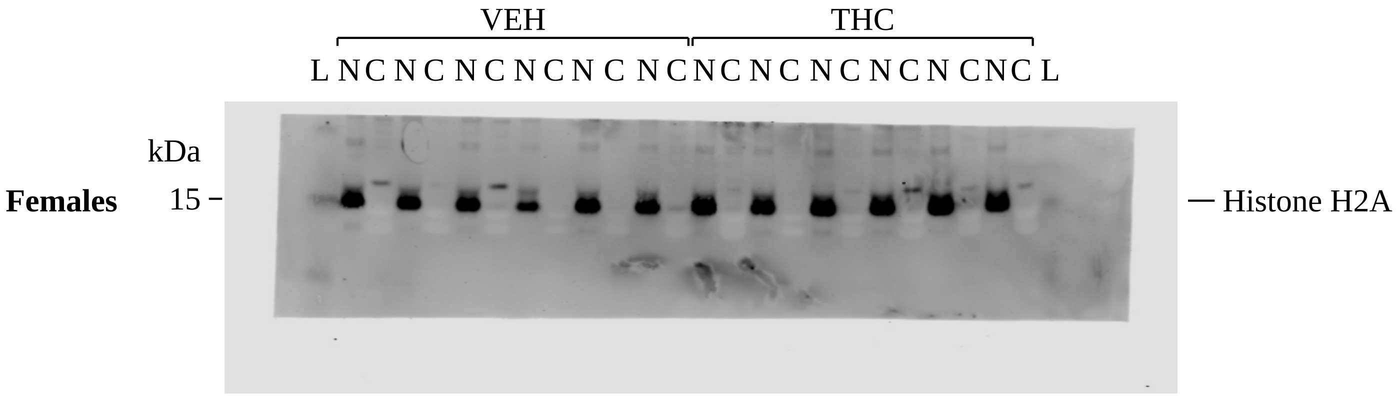


**Supplementary Figure S2.** Full-length western blots corresponding to Fig. 4. (A,B) KEAP1 and β-actin detected on the same membrane. (C) NRF2 and GAPDH in male and female placentas. (D,E) Histone H2A in male and female placentas. N, nuclear fraction; C, cytosolic fraction; L, PageRuler Plus prestained protein ladder; M, male; F, female.
